# Quantitative prediction of disinfectant tolerance in *Listeria monocytogenes* using whole genome sequencing and machine learning

**DOI:** 10.1101/2023.11.05.565740

**Authors:** Alexander Gmeiner, Mirena Ivanova, Patrick Murigu Kamau Njage, Lisbeth Truelstrup Hansen, Leonid Chindelevitch, Pimlapas Leekitcharoenphon

## Abstract

*Listeria monocytogenes* is a potentially severe disease-causing bacteria mainly transmitted through food. This pathogen is of great concern for public health and the food industry in particular. Many countries have implemented thorough regulations, and some have even set ‘zero-tolerance’ thresholds for particular food products to minimise the risk of *L. monocytogenes* outbreaks. This emphasises that proper sanitation of food processing plants is of utmost importance. Consequently in recent years, there has been an increased interest in *L. monocytogenes* tolerance to disinfectants used in the food industry. Even though many studies are focusing on laboratory quantification of *L. monocytogenes* tolerance, the possibility of predictive models remains poorly studied.

Within this study, we explore the prediction of tolerance and minimum inhibitory concentrations (MIC) using whole genome sequencing (WGS) and machine learning (ML). We used WGS data and MIC values to quaternary ammonium compound (QAC) disinfectants from 1649 *L. monocytogenes* isolates to train different ML predictors.

Our study shows promising results for predicting tolerance to QAC disinfectants using WGS and machine learning. We were able to train high-performing ML classifiers to predict tolerance with balanced accuracy scores up to 0.97±0.02. For the prediction of MIC values, we were able to train ML regressors with mean squared error as low as 0.07±0.02. We also identified several new genes related to cell wall anchor domains, plasmids, and phages, putatively associated with disinfectant tolerance in *L. monocytogenes*.

The findings of this study are a first step towards prediction of *L. monocytogenes* tolerance to QAC disinfectants used in the food industry. In the future, predictive models might be used to monitor disinfectant tolerance in food production and might support the conceptualisation of more nuanced sanitation programs.

**AUTHOR SUMMARY:** Microbial contamination challenges food safety by potentially transmitting harmful microbes such as bacteria to consumers. *Listeria monocytogenes* is an example of such a bacteria, which is primarily transmitted through food and can cause severe diseases in at-risk groups. Fortunately, strict food safety regulations and stringent cleaning protocols are in place to prevent the transmission of *Listeria monocytogenes*. However, in recent years, there has been an increase in tolerance towards disinfectants used in the food industry, which can reduce their effectiveness. In this study, we used genome sequencing and phenotypic data to train machine learning models that can accurately predict whether individual *Listeria monocytogenes* isolates are tolerant to selected disinfectants. We were able to train models that are not only able to distinguish sensitive/tolerant isolates but also can predict different degrees of tolerance to disinfectants. Further, we were able to report a set of genes that were important for the machine learning prediction and could give information about possible tolerance mechanisms. In the future, similar predictive models might be used to guide cleaning and disinfection protocols to facilitate maximum effectiveness.

## INTRODUCTION

*Listeria monocytogenes* is a concerning bacterial pathogen that is often transmitted through food and is able to cause a disease called listeriosis [1]. Infections with *L. monocytogenes* can appear sporadically or in epidemic outbreaks that sometimes span multiple countries [2]. Listeriosis is especially hazardous for at-risk groups, such as foetuses during gestation, neonates, the elderly, and immunocompromised individuals [3]. Despite having a relatively low incidence rate, the possible severity and high mortality rate of untreated *L. monocytogenes* infection underlines its importance as a foodborne pathogen and the food industry’s concern with it as a source of contamination [4,5].

Many countries have implemented strict *L. monocytogenes* regulations for food products to ensure food safety and prevent possible severe infections. For example, the USA and Turkey have adopted a “zero tolerance” policy for ready-to-eat (RTE) products, meaning that no *L. monocytogenes* is allowed to be detected in 25 g of food. Many other countries, including the EU, have adopted a risk-ranking system based on the end product, allowing low amounts of *L. monocytogenes* to be detected for RTE products where growth is not occurring [6,7]. As *L. monocytogenes* is found ubiquitously in nature, many different contamination sources exist, such as raw materials, staff, equipment, and even re-contamination within the production chain. Once in the production environment, *L. monocytogenes* isolates can persist over large timespans [8].

There are many factors contributing to the persistence of isolates, such as biofilm formation and tolerance to environmental stressors and disinfectants. Non-biological factors such as difficult-to-clean niches or improper cleaning procedures may also play a role in persistent contamination [4]. Food production sanitation protocols are thus a vital tool to prevent contamination and subsequent human infection. Quaternary ammonium compounds (QACs) constitute one of the three major disinfectant categories in the food industry [9]. The sensitivity of *L. monocytogenes* to QACs has been thoroughly studied, with several published reports of minimum inhibitory concentrations (MIC) phenotypes. However, the lack of standardised protocols and the broad variation of results due to different experimental setups make comparisons of MIC values between studies challenging [10,11].

The recent rise of interest in sequencing-based applications in the food industry and the switch of many national pathogen surveillance programs to WGS approaches gives rise to copious genome sequencing data [12]. Disentangling the complex relationships between genotypic information and phenotypic traits in hundreds to thousands of bacterial samples is a complex task. Machine Learning (ML) has remarkably succeeded in predicting genotypes and identifying genotype-phenotype relations from large datasets. For example, many studies deal with the prediction of bacterial resistance to antibiotics [13–16]. In theory, this is very similar to predicting tolerance to disinfectants. However, there is only a limited amount of studies dealing with the prediction of disinfectant tolerance from genomic data, and none of them focus on *L. monocytogenes* in a food safety context [17].

Here, we present the first and largest disinfectant tolerance prediction study for *Listeria monocytogenes* using whole genome sequencing data. Within this study, we aimed to train high-performing ML models to classify isolates into tolerant and sensitive groups for two different QAC compounds, benzalkonium chloride (BC) and didecyldimethylammonium chloride (DDAC), and one commercial disinfectant, Mida San 360 OM (MIDA). We further explored the prediction of MIC values using regression models, giving a finer-grained resolution of disinfectant sensitivity. In an attempt to account for the non-independence of the isolate data due to population structure, we implemented a phylogeny-aware nested cross-validation (CV) approach. This approach aims to prevent the predictive models from focusing on features related to phylogenetic rather than phenotypic signals. In addition, we identified a set of genomic features of importance for the prediction of disinfectant tolerance by our ML models that might indicate possible genotype-phenotype relationships. Lastly, to assess the generalisation performance of our models to previously unseen data, we evaluated our pre-trained models on three independent test sets. The results from this study show great potential for the prediction of disinfectant tolerance from genomic data. In the future, our predictive models might aid the food industry in designing strategic disinfection plans and could help guide targeted elimination in contamination situations.

## MATERIALS AND METHODS

### Data acquisition

The data for this study was extracted from a large phenotypic and genomic characterisation study of *Listeria monocytogenes* by Ivanova et al. [10]. Within that study, 1,671 *L. monocytogenes* isolates were tested for their sensitivity to QACs. In particular, the study reported disinfectant tolerance phenotypes for two pure substances (benzalkonium chloride and didecyldimethylammonium chloride) and one commercial disinfectant (Mida San 360 OM). In addition, they shared WGS data as well as metadata for all isolates.

For our study, we are using a subset of 1649 isolates from the study. We excluded all samples from *L. monocytogenes* lineages III and IV (n=18), as all the samples from these lineages are sensitive to BC. Two more samples were excluded because of inconsistencies in file names and raw read headers, which raised issues during mapping, and one additional sample was excluded due to low depth of coverage. Additionally, we excluded one isolate whose assembly genome size was higher than expected (> 3 Mb ± 0.5 Mb), which could indicate contamination or sequencing quality issues. A list of isolate IDs, accession numbers, metadata, and disinfectant MICs can be found in Table S1.

Whereas all the isolates were laboratory tested for BC sensitivity, only a subset of isolates were tested for DDAC (n=247) and MIDA (n=154). Apart from the minimum inhibitory concentration (MIC) values, we also extracted the thresholds to classify the isolates into tolerant and sensitive. These thresholds are ≥ 1.25 mg/L, > 0.4 mg/L, and ≥ 0.63 mg/L for BC, DDAC, and MIDA, respectively.

### Sample pre-processing and feature extraction

In this study, we compared two different genetic inputs, i.e., single nucleotide polymorphisms (SNP) and pan-genome gene cluster features, as inputs to the ML prediction algorithms. We extracted the SNP features from the raw read sequencing data using the workflow described in the S1 Appendix. To extract the gene cluster features, we first assembled all genomes using SPAdes (v3.11.0) [18] and then screened all assemblies for the presence or absence of the pan-genome gene clusters, as described in the S2 Appendix. The screening reports a percent identity measure for each reference pan-genome gene cluster, which is subsequently used as input for machine learning.

### Machine Learning

The input for our machine learning pipeline consists of the extracted genomic feature data (i.e., SNPs or pan-genome gene clusters) and the laboratory phenotype data. For classification, the isolates were separated into a tolerant and sensitive class using the thresholds of ≥ 1.25 mg/L, > 0.4 mg/L, and ≥ 0.63 mg/L for BC, DDAC, and MIDA, respectively. To assess if we could get a finer-grained prediction of biocide tolerance, we used regression models to predict MIC values directly. MIC values are usually reported as discrete concentration values based on a two-fold dilution series (i.e., logarithmic scale), making evaluating errors less intuitive. Hence, we also tested the prediction ability on log_2_ transformed MIC values, making evaluating errors more intuitive and removing scale-dependent effects.

Choosing a ML model that is well suited to capture the underlying relationships of features and outcome is not straightforward. To optimise time and save computational resources, we pre-screened various ML models for their predictive performance using the BenchmarkDR pipeline (https://github.com/WGS-TB/BenchmarkDR) [19]. BenchmarkDR is a modular pipeline that uses genomic short-read information to benchmark (i.e., systematically compare) different models that can predict phenotypes. Even though BenchmarkDR can also benchmark the pre-processing and extraction of genomic features, we only used it for the model training part. In short, BenchmarkDR performs a repeated (n=5) stratified shuffle split with an 80%/20% training/test split using the sci-kit learn library (v1.2.1) [20] in Python (v3.8.13) and reports model performance with various measures. The model classes with the best performance identified by BenchmarkDR were used in the final training.

The assumption of sample independence plays an important role for many ML models. However, bacterial population structure (i.e., genomic relatedness or phylogenetic proximity) creates underlying dependencies between isolates [21,22]. To mitigate this, we used grouped *k*-fold cross-validation with phylogenetic information. In this “phylogenetically aware” ML, phylogenetically close isolates are kept together as a group, which is not split between the training and testing set. This should reduce overfitting and overly optimistic prediction performance from simultaneously having similar isolates in the training and testing set [23,24]. We compared three different methods to group the samples phylogenetically. These were the well-established concepts of clonal complexes (CC), k-mer clustering using kma (v1.3.15) [25] with a k-mer identity cut-off of 95%, and variable-length k-mer clustering using popPUNK (v2.5.0) [26].

The model training was performed with nested cross-validation for BC tolerance classification and regression. The outer grouped *k*-fold CV was realised with k=10. The inner grouped *k*-fold (k=5) was used to tune the models’ hyperparameters with Bayesian optimisation for 30 iterations using the scikit-optimize library (v0.9.0) [27]. The cross-validation performance of the individual models was reported with balanced accuracy (BA) scores and mean squared error (MSE) for classification and regression, respectively.

Due to the limited amount of isolates tested for DDAC and MIDA, we only performed classification and omitted MIC prediction via regression. Additionally, we changed the grouped *k*-fold parameter to k=5 and k=3 for outer CV and inner CV, respectively, and only used CC cluster information to account for phylogenetic relatedness. The code for all our ML models can be found on GitHub (https://github.com/agmei/LmonoDisinfectML).

### Independent test sets

To evaluate how well our ML predictors generalise to unseen data, we used an independent set of isolates not included in the model training process. We extracted data from three studies that shared WGS data to public repositories and reported BC MIC values or categorical phenotype information (i.e., sensitivity/tolerance) [28–30]. The genome sequencing accession numbers and metadata of the 286 extracted isolates can be found in Table S2. We excluded nine out of 197 isolates from Palma et al. [29]. We could not access the genomic data on the public repository for three of these nine isolates. For the other six, we were not able to extract MIC values due to the discordance of isolate IDs in the metadata. We extracted the MIC values/phenotypes of the remaining isolates and downloaded the corresponding WGS data. To be able to compare the classification predictions from our trained model with the validation set, we categorised the MIC values into tolerant and sensitive groups (i.e., phenotypes). This has been done as described by Ivanova et al. [10]. In short, for the publications that do not report an MIC cut-off for tolerant/sensitive separation, we screened all isolates for the presence of one of four genes associated with QAC resistance (qacH (multiple variants), bcrABC (JX023284.1), emrE (NC_013766.2:c1850670-1850347), and emrC (MT912503.1:2384-2770)). The threshold was then set at the lowest MIC value for which at least one of the four genes was present, where the presence of a gene is defined by a sequence identity of ≥ 90% at the nucleotide level. The results of the alignment, as well as the resulting phenotype categories, can be found in Table S2. A more detailed description of the feature extraction from the validation dataset can be found in the S3 Appendix.

### Feature importance with SHAP

We used Shapley additive explanations (SHAP) values to evaluate which features drive our ML in decision-making [31,32]. To make the explanations more robust and get an understanding of the feature importance using the entire dataset, we cross-validated our SHAP calculation using *5*-fold CV. As a result, we get SHAP values that have been evaluated for each isolate [33]. In order to get insight into possible functions of the important features (i.e., pan-genome gene clusters), we manually annotated the ten most highly ranked features (ranked according to their SHAP value) using complete reference genomes from NCBI’s RefSeq database [34]. See the S4 Appendix for a detailed description of the manual annotation.

## RESULTS

### Pan-genome features outperform SNP features for BC

To see the effect of genomic input on the ML prediction, we compared the predictive performance of several ML models using pan-genome features (n=14531) or SNP features (n=316084) as input. For classification into tolerant and sensitive classes, using pan-genome features resulted in higher cross-validated BA scores for 13 out of 14 models (Table S3). Only for logistic regression with stochastic gradient descent and L2 regularisation (SGDC) did using SNP features result in a higher BA score (0.87) than using pan-genome features (0.54). A similar pattern appears for the regression tasks, i.e., prediction of MIC and log_2_ transformed MIC values. For both tasks, five out of seven models resulted in a lower mean squared error when trained on pan-genome features (Table S3).

### Pre-screening of selected tree-based and linear ML models for further evaluation

As the relationship between genotype and phenotype can be quite complex, it is difficult to foresee what kind of ML model is most suited to capture the underlying signal. Hence, we tested a broad range of ML models and compared their CV performance as part of a pre-selection process. Looking at the results for classification (Fig 1a), we see that many of the models performed similarly, reaching BA scores of up to 0.98 for the gradient-boosted tree classifier.

**Fig 1.**
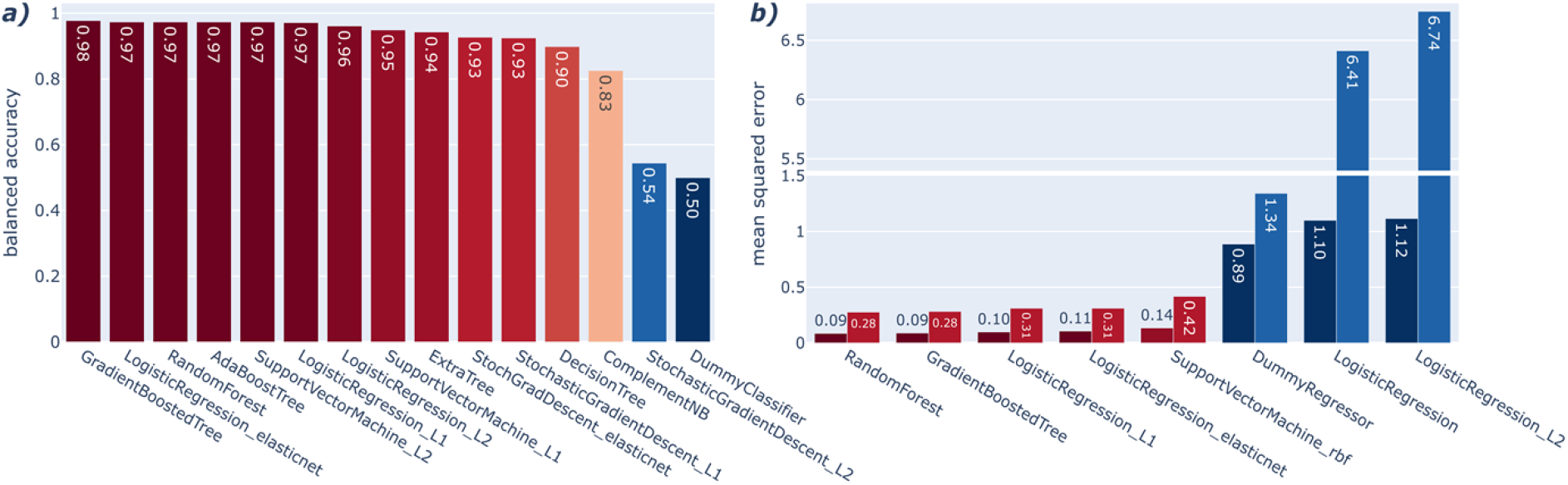
Pre-screening a variety of ML models with BenchmarkDR. **a)** Classification performance comparison of the cross-validated balanced accuracy results from the pre-screening. The colour scale describes the model’s performance from red (good) to blue (poor). **b)** Regression performance comparison of the cross-validated mean squared error results from the pre-screening. The colour scale describes the model’s performance from red (good) to blue (poor). The darker shades (right columns) correspond to the regression of non-transformed MIC values, and the lighter shades (left columns) correspond to the regression of log_2_ transformed MIC values.

The results of the regression tasks (Fig 1b) show that we were able to train well-performing models reaching mean squared errors (MSEs) of 0.09 and 0.28 for regression of the untransformed and log_2_ transformed MIC values, respectively. For both regression tasks, the best-performing models are random forest and gradient-boosted tree regressors. We also see that linear regression and linear regression with L2 regularisation perform worse than a dummy regressor that predicts the median of the test set.

### Phylogeny-aware ML reported similar performances across the compared clustering methods but vastly different performances between models

To account for sample relatedness, we compared the effect of three different clustering methods (i.e., CC, kmer95, and popPUNK clusters) on the prediction performance. For this analysis, we used three ML models (random forest (RF), gradient-boosted trees (GBT), and logistic regression with L1 regularisation (LR_L1)), which have been selected in the pre-selection screening.

A comparison of the clustering methods by normalised mutual information [35] showed high concordance between clustering with scores of 0.98, 0.94, and 0.93 for CC-kmer95, kmer95-popPUNK, and CC-popPUNK, respectively.

The classification results only showed a slight difference in performance across clustering methods (Fig 2a). In particular, for RF and GBT models, the medians and interquartile ranges (IQR) overlap indicates that these grouping methods are similar. For LR_L1, we can see that the median for kmer95 clustering is slightly above the IQR of the other two clustering methods. However, the mean for this method is within one standard deviation of the other two methods (Table S4).

**Fig 2.**
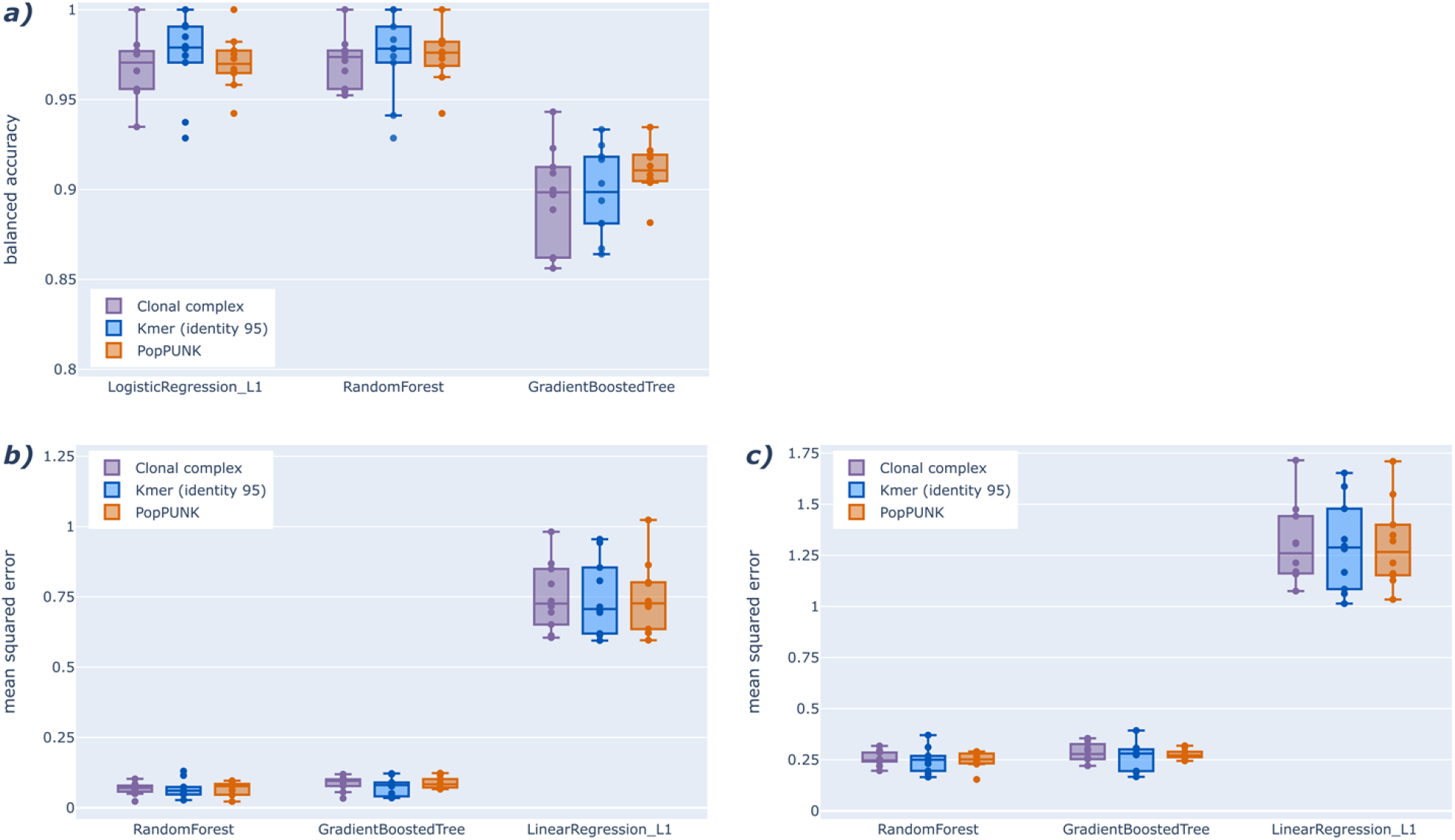
Comparison of three different clustering methods for phylogeny aware ML model training. **a)** Cross-validated classification performance scores for pre-selected models and different phylogenetic clustering methods. **b)** Cross-validated regression performance for the pre-selected models and different clustering methods. **c)** Cross-validated regression performance for log_2_ transformed MIC values of the pre-selected models across the different clustering methods.

The MSE results for the regression tasks show a similar pattern (Fig 2a and 2b), as all mean performance scores are within one standard deviation of each other (Table S4).

Comparing the classification performance between the different models shows a better performance (i.e., higher BA) for the LR_L1 and RF models than for GBT models. On both regression tasks, we observed a better performance (i.e., lower MSE) for the RF and GBT models than for linear regression with L1 regularisation (LinR_L1).

### Feature importance analysis identified pan-genome features driving ML predictions

In order to better understand the importance of specific features for the ML models’ phenotype prediction, we conducted a post-hoc feature importance analysis with SHAP. In particular, we used the CC-aware linear models with L1 regularisation and random forest models for the classification and regression tasks, respectively. As some of our pan-genome gene clusters were missing annotation, we screened the top ten most important features by absolute mean SHAP value against a non-redundant set of annotations extracted from complete reference genomes. The complete list of the pan-genome features and their SHAP values can be found in Table S5.

For classification, only two out of the top-scoring features have an annotation, namely *qacC*, a small multidrug resistance (SMR) efflux transporter, and *ebrB,* a SMR gene (Fig 3a). The reported pan-genome gene cluster names are based on Prokka’s reference databases and might not be fully concordant with species-specific naming, e.g., *qacC* would correspond to *qacH* in *L. monocytogenes*. The other eight are ‘hypothetical gene clusters’ that were assigned a group number during the pan-genome analysis using Roary. Our analyses identified the pan-genome gene cluster group 6155 (*qacH)* to have the most importance to the ML prediction. The manual annotation resulted in eight out of ten matches with protein sequence identities of > 90% (Table 2 and S5). These matches are comprised of hits to transcriptional regulators, efflux transporters, transposases, cell wall anchoring domains, and phage-related proteins. For the remaining features, the search resulted in matches with identities in the 60-80% range, matching with TetR transcription regulator and phage proteins.

**Fig 3.**
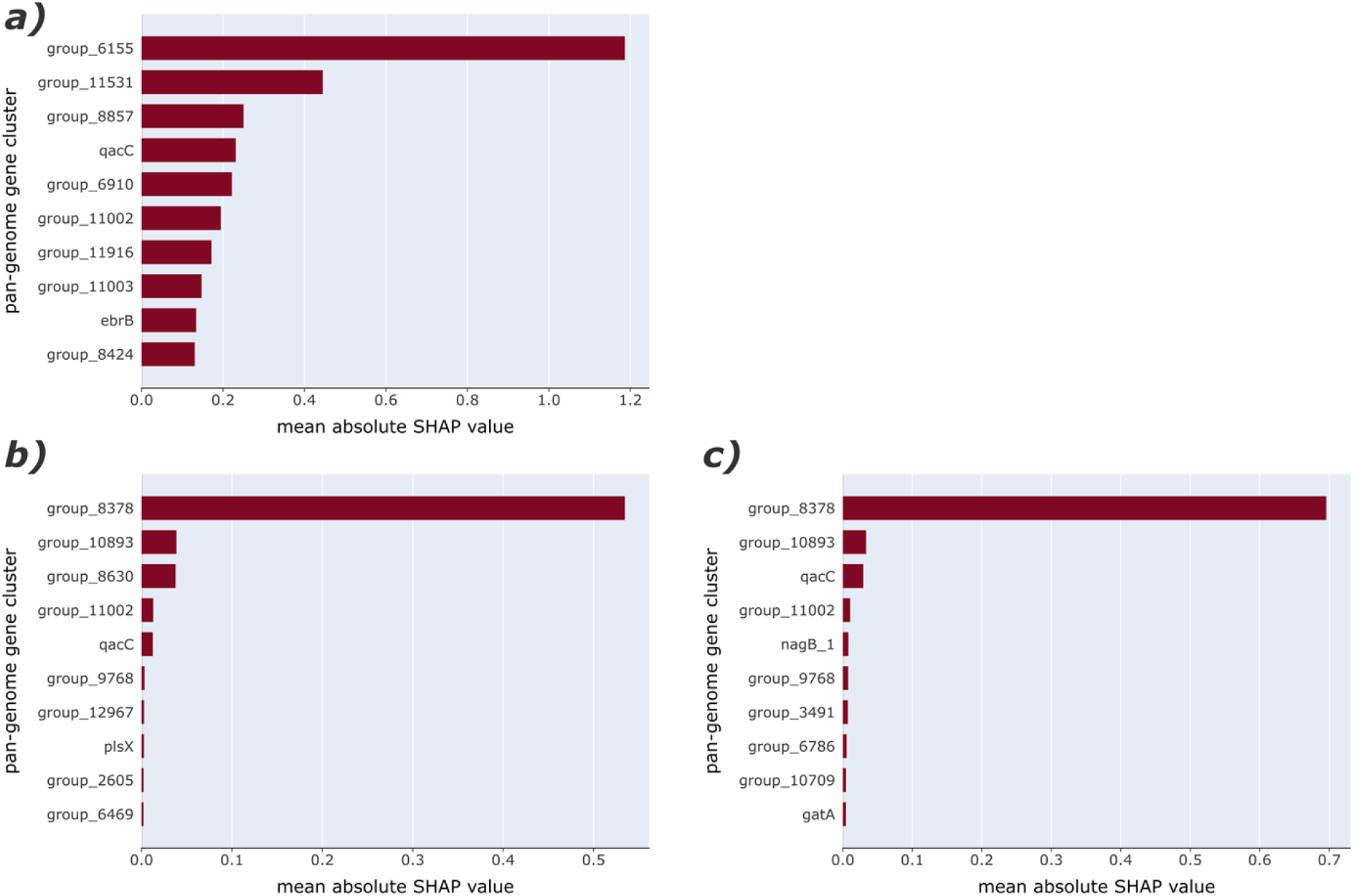
Feature importance evaluation using SHAP values and CC clustering. **a)** Top ten most important features for the logistic regression with L1 regularisation into sensitive/tolerant categories. **b)** Ten most important features for the random forest regression of MIC values. **c)** Ten most important features for the random forest regression using log_2_ transformed MIC values.

**Table 2.** Manual annotation results for the top ten most important classification features.

| Pan-genome gene cluster | Protein Description | Identity (%) | Coverage (%) | E-value |
| --- | --- | --- | --- | --- |
| **group_6155** | TetR family transcriptional regulator | 100 | 96 | 1,2E-95 |
| **group_11531** | LPXTG cell wall anchor domain-containing protein | 100 | 13 | 3,2E-07 |
| **group_8857** | DUF6262 family protein (TnpC) | 100 | 99 | 1,0E-75 |
| ***qacC*** | Quaternary ammonium compound efflux SMR transporter QacH | 100 | 99 | 6,0E-81 |
| **group_6910** | HK97 gp10 family phage protein | 100 | 100 | 1,4E-83 |
| **group_11002** | Quaternary ammonium compound efflux SMR transporter QacH | 92,6 | 97 | 6,5E-70 |
| **group_11916** | DUF1642 domain-containing protein | 60,6 | 57 | 2,1E-54 |
| **group_11003** | TetR family transcriptional regulator | 76,7 | 40 | 1,8E-69 |
| *ebrB* | Quaternary ammonium compound efflux<br>SMR transporter BcrC | 100 | 100 | 3,6E-76 |
| **group_8424** | hypothetical protein (RepC) | 100 | 99 | 3,9E-66 |

Only two important features for the regression of MIC values already have an annotation, namely *qacH* and *plsX*, the latter involved in fatty acid/phospholipid synthesis (Fig 3b). For regression of log_2_ transformed MIC, there are three other annotated features: *qacC*, *nagB_1*, a cell-wall peptidoglycan gene, and *gatA*, a glutamyl-tRNA(Gln) amidotransferase (Fig 3c). In both cases, group 8378 (*emrE*) and group 10893 (*qacH*) have been identified as the top two most important features, with group 8378 being far more important in comparison to all others (Table S6).

### Evaluation of independent test sets shows performance benefits for linear models

To evaluate how well our classification models perform on independent test datasets, we collected data from three studies that reported BC phenotypes and shared WGS data for their isolates. Here, we present the findings for phylogeny-aware ML with CC clusters (Table 3). Overall, the balanced accuracy scores range from 0.5 for the GBT classifier to 0.67 for the LR_L1 classifier. Looking at the data from the different studies separately, we see large performance differences, with the highest BA of 0.93, 0.78, and 0.61 for data from Cooper et al. (2021), Cherifi et al. (2018), and Palma et al. (2022), respectively. There are considerable differences between the three compared models, both overall and for the individual datasets. LR_L1 yielded the highest performance for Cooper et al., Cherifi et al., and overall. The RF classifier performed best for Palma et al.. The GBT classifier performed the worst for all datasets, consistently yielding a BA of 0.5, which indicates a prediction performance that is no better than random. Prediction performance for the other phylogenetic clustering methods showed similar results, with LR_L1 performing best overall and when assessing the datasets individually (Table S7).

**Table 3.**
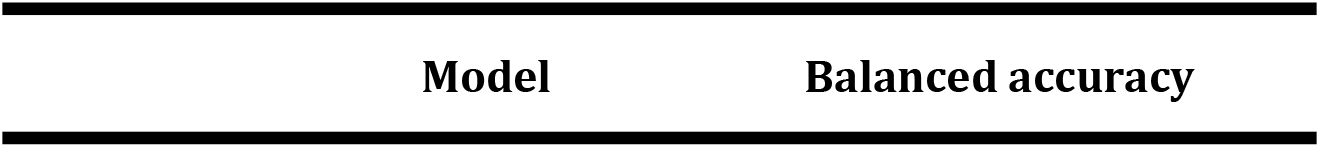
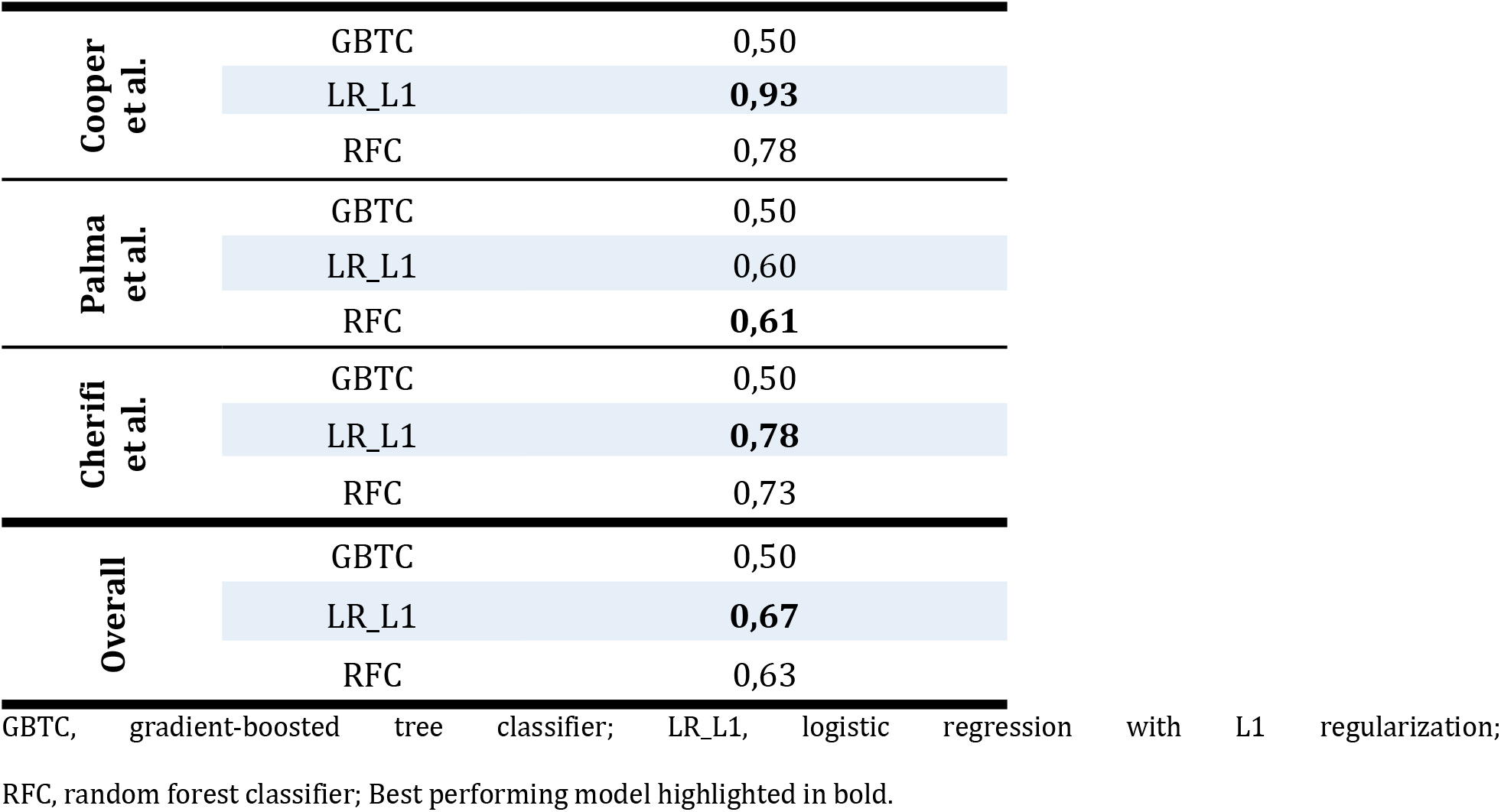
Performance evaluation for phylogeny-aware ML with CC clusters on the independent test sets.

| Model | Balanced accuracy |
| --- | --- |

|  |  |  |
| --- | --- | --- |
| Cooper<br>et al. | GBTC | 0,50 |
|  | LR_L1 | **0,93** |
|  | RFC | 0,78 |
| Palma<br>et al. | GBTC | 0,50 |
|  | LR_L1 | 0,60 |
|  | RFC | **0,61** |
| Cherifi<br>et al. | GBTC | 0,50 |
|  | LR_L1 | **0,78** |
|  | RFC | 0,73 |
| Overall | GBTC | 0,50 |
|  | LR_L1 | **0,67** |
|  | RFC | 0,63 |

GBTC, gradient-boosted tree classifier; LR\_L1, logistic regression with L1 regularization; RFC, random forest classifier; Best performing model highlighted in bold.

### The predictions for other QAC disinfectants (DDAC and MIDA) show promising results

Besides BC, DDAC is another commonly used QAC disinfectant in the food industry. It is also one of the active compounds in the commercial disinfectant MIDA. To see how well we can predict phenotypes for other QAC disinfectants, we used the subset of isolates for which disinfectant sensitivity to DDAC and MIDA has been tested and trained predictive ML models on them. Similarly to BC, we pre-screened different models with BenchmarkDR and trained phylogeny-aware ML models of the selected types. In contrast to the analysis of BC, we only used CC clusters phylogeny-aware ML, as we did not observe major differences between the clustering methods in the BC analysis. The pre-screening results show that many ML models perform similarly well (Fig 4a&b). We choose five models: logistic regression with L1 regularisation, random forest, gradient-boosted trees, AdaBoost trees (ABT), and support vector machines with L1 regularisation (SVM_L1). The models were chosen based on their performance and ability to explore linear and non-linear relationships in the data. Nested-CV results for DDAC in the phylogeny-aware models showed that the LR_L1, ABT, and SVM_L1 perform similarly, as indicated by the overlapping medians and IQRs as well as all means being within one standard deviation of each other. The RF and GBT models perform worse than the others (Fig 4c and Table S4). The nested-CV results for MIDA show a similar trend (Fig 4d and Table S4).

**Fig 4.**
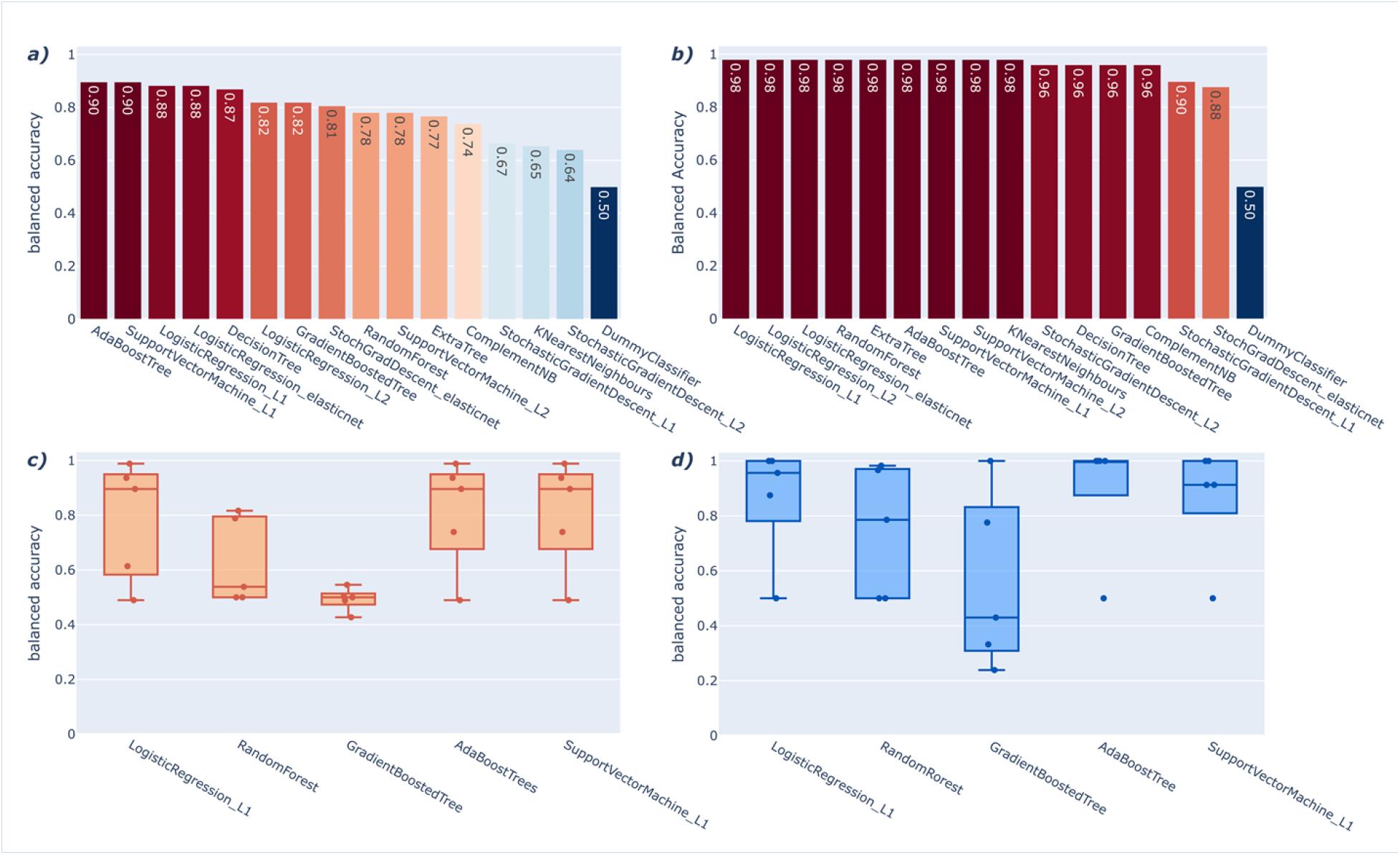
Cross-validated ML performances for alternative QAC compound DDAC and commercial disinfectant MIDA. **a)** DDAC pre-screening BA results using BenchmarkDR. **b)** MIDA pre-screening BA results using BenchmarkDR. **c)** Phylogenetically aware CV BA performance for DDAC using pre-selected models and CC as clustering. **d)** Phylogenetically aware CV BA performance for MIDA using pre-selected models and CC as clustering.

In order to assess whether ML models trained on pure compound data are indicative of tolerance to commercialised products, we used our pre-trained DDAC models to predict phenotypes of isolates tested for MIDA, as it contains DDAC as one of its main active agents. The results show remarkable concordance of the DDAC model predictions with the MIDA phenotypes, reaching balanced accuracy scores of 0.96 for the L1 logistic regression and AdaBoost models (Table S4).

## DISCUSSION

Many bacteria tend to have strong population structures, which results in non-independence between data points and can have substantial implications on the generalisation to new data and the importance of features for prediction. In particular, this could select/highlight features related to the phylogenetic background signal (lineage effects) rather than the phenotype. While this phenomenon might not affect the training dataset performance, it could lead to poor predictive performance on new datasets with different phylogenetic distributions. There is currently no consensus on how to address these problems. Some studies account for population structure during model training by keeping closely related groups separate during cross-validation (i.e., by not letting them get split between training and testing sets) [23,24]. This method attempts to limit machine learning from learning isolate predictive features, as phenotypic traits often correlate with isolate structures [23]. This method has been applied to different ML problems with structured data and is regarded as ‘blocking’ [21,36,37]. We used a grouped K-fold split to account for population structure during training.

Additionally, we were comparing the effect of different phylogenetic structure clustering methods on the predictive performance. Our results do not indicate any significant difference between the methods. Hence, we use CC structure models for our target application as they are easily implementable and a well-established method to cluster *L. monocytogenes* isolates, for example, in epidemiological surveillance.

Another critical aspect of performance evaluation for ML models is their generalisation ability to previously unseen data (i.e., independent test sets). We tested our pre-trained ML classifiers on laboratory-confirmed BC sensitivity phenotypes collected from three studies. Overall, our models performed modestly and vastly underperformed our nested-CV performance estimation. Taking a closer look at the datasets individually, we found considerable variation in prediction performance. In particular, for the Palma et al. data, performance scores peaked at 0.61, a more than 30% difference from the training performance of BA 0.97.

In general, comparing results from disinfectant sensitivity testing experiments is quite challenging, as the MIC varies greatly depending on experimental design, such as biocide and inoculum concentrations and cultivation media, temperature, and time [10]. This could potentially affect the generalisation ability of the tested ML models. Another factor that can result in lower prediction performance than expected is if the new dataset includes different sub-populations than the training set. However, looking at the metadata of the independent test set (Table S2), this seems not to be an issue in this case.

Further investigation described in the S5 Appendix showed an almost perfect concordance of the predictions with the absence/presence of QAC genes. The ML classifier classified all isolates with QAC genes as tolerant and all isolates without QAC genes as sensitive, except for one isolate. This may explain why the prediction performance for the Palma et al. dataset is so much lower than expected. If we were to adjust the separation criteria to having or not having QAC genes present for sensitive and tolerant classes, respectively, we would raise the balance accuracy performance from 0.60 to 0.97. These findings lead us to believe that the way we categorise the independent test set isolates might not be optimal and results in limited validation strength for our trained ML models.

After screening a diverse set of ML models for their ability to predict tolerance to benzalkonium chloride, we have to assess which model should be used for predictions in practice. The results for the classification of isolates lead us to conclude that logistic regression with L1 regularisation performs best, with a mean cross-validated balanced accuracy performance of 0.97±0.02. Even though we could not see any apparent performance differences for LR_L1 in the phylogeny-aware ML, we observed an advantage in using this model on the independent test sets, where it had the highest performance for two of the three datasets and overall.

Our analysis showed promising results for direct MIC prediction, and we identified RF and GBT models as top performers with cross-validated MSE scores on log_2_ transformed MIC values of 0.25±0.03 and 0.28±0.05, respectively. Even though only a limited number of studies have explored MIC prediction of disinfectants from genomic data, making comparisons difficult, tree-based models show good performance and are commonly used for similar prediction tasks [15,16,38,39].

Understanding how a ML model arrives at its prediction can give valuable insights into the underlying patterns or processes. Even though one has to be cautious in understanding that correlational signals do not equal causal relations, uncovering patterns between features and outcome variables could be indicative of biological relations. Analysing our final model for the classification of BC tolerance (Logistic Regression with L1 regularisation and CC phylogenetic clustering) using SHAP values, we were able to rank all features according to their importance. Unfortunately, many of the pan-genome gene cluster features lack annotation.

To get a better understanding of possible functions for some of the unannotated features, we manually annotated the ten highest-ranked features. Three out of the ten features resulted in hits for well known QAC resistance mechanism in *L. monocytogenes*, i.e., QAC efflux small multidrug resistance transporter QacH or BcrC. Both mechanisms have previously been linked to enhanced QAC tolerance [40,41]. The *bcrC gene,* which is part of the *bcrABC* cassette, was initially annotated by Prokka as *ebrB,* which was most likely a misannotation due to a lack of species-specific naming in annotation databases. Two features resulted in hits for TetR family transcriptional regulators. Such regulators are a known part of the *bcrABC* cassette and regulate the *emrC* and *qacH* genes, which are all associated with BC tolerance [4,42]. In addition, Kropac et al. found that BC exposure strongly induces the expression of a TetR transcription regulator in the pLMST6 plasmid, which is known to carry the *emrC* gene [43]. We further found two features with possible association to plasmids. One matched the *tnpC* gene, encoding a transposase which is part of the Tn*6188* transposon and has been found to confer BC tolerance, and the other one resulted in a match for *repC,* which codes for proteins involved in plasmid replication [40,44]. Another two features returned hits for phage-related proteins. Phages are known to play a role in the horizontal transfer of genes, such as antimicrobial resistance genes [45]. One feature resulted in a hit for an LPXTG-domain-containing protein, which is a part of cell wall anchors but has no known link to disinfectant tolerance thus far. For the regression tasks (i.e. MIC prediction and log_2_ MIC prediction), we analysed the random forest models with CC phylogenetic clustering. We found that the top two most important features are the same for both tasks, with group 8378 having the biggest influence on the prediction. Manual annotation resulted in matches for EmrE and QacH multidrug transporters, both known mechanisms for QAC tolerance in *L. monocytogenes* [46].

Extending our methodology to different compounds used in the food industry (DDAC) as well as a commercial disinfectant (MIDA) is broadening our view on applicability in a production setting. Similar to BC, we were able to train well-performing ML classifiers for both DDAC and MIDA, reaching balanced accuracy scores of up to 0.81±0.18 and 0.90±0.20, respectively. However, we find that for both disinfectants, the prediction performance varies greatly between different CV-splits, which indicates limited model robustness. This might be due to the small amount of data points (only ∼150-250 isolates) used for the training, making it hard for the ML to focus on patterns that generalise. More studies with increased isolate data would be beneficial to ensure a trained ML model with good robustness for application in the industry.

In practice, the implementation of genome sequencing applications in the food industry is relatively slow. Imanian et al. have described some essential factors that influence the food industry’s interest in high-throughput sequencing, such as improvement or verification of control measure design, reduced food safety management costs, and the increased shift of governmental agencies towards WGS-based applications [12]. Our study proposes another possible application of WGS that could improve *L. monocytogenes* control measure design and reduce food safety management costs. Predictive ML models, in combination with WGS, could be used in the future to internally monitor disinfectant tolerance in production plants and assess the efficacy of sanitation measures.

Recent studies have observed great concordance (up to 99%) between known QAC resistance genes and tolerant/sensitive phenotypes, raising the question of the benefit of a more complex ML solution [10]. In this dataset, 20 isolates showed tolerance to BC in laboratory experiments, but no QAC resistance genes were observed. Our L1 logistic regression model classified 8 of these 20 isolates as tolerant. This indicates a possible benefit of using ML over a classification with known QAC genes alone. An additional benefit of our proposed method is that we are not limiting our prediction to only a small set of known genes so that we may discover relationships between a more extensive set of genes and the phenotype.

Even though our study shows promising results for predicting disinfectant tolerance using ML, some general limitations could influence the transferability of the results from our method to a food industry setting. In practice, disinfectants are rarely used as pure compounds but are part of commercialised products combining different active agents. We have already tried to address this limitation by including a commercial disinfectant, Mida San 360 OM, in our analysis. Additionally, we evaluated whether the results from pure compound studies are indicative of tolerance to commercialised products using our pre-trained DDAC models to predict MIDA phenotypes. The analysis showed promising results indicating the possible transferability of prediction models trained on pure compounds to commercialised disinfectants.

Another major limitation is that our study relies on phenotypic data of single species cultures. Many studies have found that *L. monocytogenes* tends to form biofilms in food production settings, making proper eradication more complex [47–49]. Thus, it will be crucial for future models to incorporate biofilm factors to obtain predictive models that better simulate the application in practice.

In general, many factors, such as the additional compounds and host-specific interactions, may play a role in the disinfection process. Hence, more research is needed to better understand the transferability of pure compound models to actual industry settings.

## CONCLUSION

We were not only able to obtain high-performing ML classifiers for the prediction of disinfectant sensitivity/tolerance from WGS data but also to show promising results for the prediction of MIC values directly, resulting in a more detailed phenotype separation. Additionally, we reported important features steering the ML prediction that could be indicative of possible genotype-phenotype relationships and could be used as a starting point for laboratory confirmation studies.

As WGS analysis finds more popularity in the food industry, there is a great variety of application possibilities. For example, predictive models like the one proposed in this study might be beneficial to monitor disinfectant tolerance in food production sites and to aid the development of more effective sanitation programs. In addition, this study serves as a proof-of-concept for predicting disinfectant tolerance using WGS and ML. This methodology could be extended to other disinfectant agents, such as peracetic acid, for which tolerance mechanisms are mostly unknown.

However, there are still some limitations that minimise the transferability of the current models to an industry setting. In practice, many factors, such as biofilm formation, mediate the survival of sensitive *L. monocytogenes* isolates that are not considered in this study. Nevertheless, this study is an essential first step towards predicting disinfectant tolerance from WGS data for application in a food industry setting and sets priority directions for further research.

## ACKNOWLEDGEMENT

PL received support for this work by Karl Pedersen og Hustrus Industrifond (DI-2019-07020), the Danish Dairy Research Foundation, and the Milk Levy Fund. LC acknowledges funding from the MRC Centre for Global Infectious Disease Analysis (reference MR/X020258/1), funded by the UK Medical Research Council (MRC). This UK funded award is carried out in the frame of the Global Health EDCTP3 Joint Undertaking.

## SUPPORTING INFORMATION

See attached file.

